# A common mechanism of temperature-sensing in thermoTRP channels

**DOI:** 10.1101/2022.05.23.493163

**Authors:** Lizhen Xu, Xiao Liang, Wenxuan Zhen, Zhangzhi Xue, Fangfei Zhang, Xiao Yi, Xiaoying Chen, Lidan Hu, Bei Li, Bing Zhang, Yuhang Wang, Zhenfeng Deng, Wei Yang, Han Wen, Tiannan Guo, Yi Zhu, Fan Yang

**Author notes:** These authors contributed equally to this work (Lizhen Xu, Xiao Liang, Wenxuan Zhen). Correspondence should be addressed to: Fan Yang, Yi Zhu, Tiannan Guo, or Han Wen.

## Abstract

Detecting temperature is crucial for the survival of living organisms. Though the thermo transient receptor potential (thermoTRP) channels, such as TRPV1 or TRPM8, have been identified as prototypic heat or cold sensors, respectively, how they detect temperature remains elusive. Here we first identified groups of clustered residues in these channels that undergo burial/exposure conformational rearrangements during temperature activation by analyzing available protein structures or hydroxyl radical footprinting-mass spectroscopy (HRF-MS). By systematically perturbing water-protein interactions at these residues, we found that the temperature sensitivity in these channels were modulated in accordance with the sidechain hydrophobicity. The changes in energy associated with changes in water-protein interactions were sufficient for thermo activation. Therefore, our study has established that the water-protein interactions as a common mechanism underlying temperature sensing in TRPM8 and TRPV1.

## INTRODUCTION

All atoms of proteins in living creatures are in constant thermal motion. When temperature is changed, the thermal motion and the activities of a protein will change as well. However, there are a group of thermo-sensitive TRP channels (thermoTRPs), exemplified by the heat sensor TRPV1(Caterina et al., 1997) and the cold sensor TRPM8(McKemy et al., 2002; Peier et al., 2002), whose open probability is exquisitely sensitive to temperature. For instance, when TRPV1 channel is opened by noxious heat (> 42℃) or TRPM8 channel is activated by low temperature (< 28℃), their temperature dependence of current increase as calculated by Q10 is well above 20 and their enthalpic and entropic changes in temperature gating are much larger than those of regular proteins(Julius, 2013; Zheng, 2013). Moreover, these thermoTRPs serve as temperature detectors *in vivo*, so when they are genetically knocked out(Caterina et al., 2000; Dhaka et al., 2007) or pharmacologically inhibited(Moran, 2018), the temperature sensing ability in animals was largely disrupted. Despite the critical physiological roles of thermoTRPs, how these channels detect temperature and whether there is a common temperature sensing mechanism employed in heat or cold-sensing TRP channels remain largely elusive.

We proposed a hypothesis that changes in the burial/exposure state (therefore the water-protein interactions) of specific amino acids as the mechanism of temperature sensing in thermoTRPs. For instance, when temperature is decreased and the thermo motion of water molecules is reduced, a hydrophobic sidechain of TRPM8 residue should be stabilized in the exposed state. If the hydrophobicity of such a residue is increased, the mutant TRPM8 is expected to be more cold-sensitive. For TRPV1, if a residue becomes more exposed in heat activation, increasing sidechain hydrophilicity would enhance heat sensitivity as opposite to TRPM8. In the voltage-gated potassium channel, altering the hydrophobicity of exposed residue changes heat sensitivity of the channel (Chowdhury et al., 2014). A theoretical analysis suggested that the temperature sensing apparatus in TRP channels could be distributed on a single-residue basis, where changes in the burial/exposure state of 10-to-20 residues per subunit would be enough to drive temperature activation (Clapham and Miller, 2011). Though previously we have employed the fluorescent unnatural amino acid 3-(6-acetylnaphthalen-2-ylamino)-2-aminopropanoic acid (ANAP) to identify three residues in TRPM8 with buried to exposed conformational changes upon cooling and preliminarily tested our hypothesis (Yang et al., 2020), there are obvious limitations. For instance, the bulky sidechain of ANAP often caused the channel with ANAP incorporation to be unfunctional, leading to missed identification of residues. Therefore, our water-protein interaction hypothesis of temperature sensing remains to be tested with more extensive identification of residues with burial/exposure changes in thermoTRPs.

To rigorously test our hypothesis of temperature sensing, we focused on the prototypic heat sensor TRPV1 and cold sensor TRPM8. There were two tasks involved in testing our hypothesis, the first was detecting residues with conformational changes during temperature activation, and the second was functional validation by measuring temperature sensitivity in mutants with altered sidechain hydrophobicity (SCH) (Figure 1A). We employed solvent accessible surface area (SASA) measurements from cryo-EM structure models and HRF-MS in TRPV1 and TRPM8, respectively, for the first task as detailed below.

**Figure 1.**
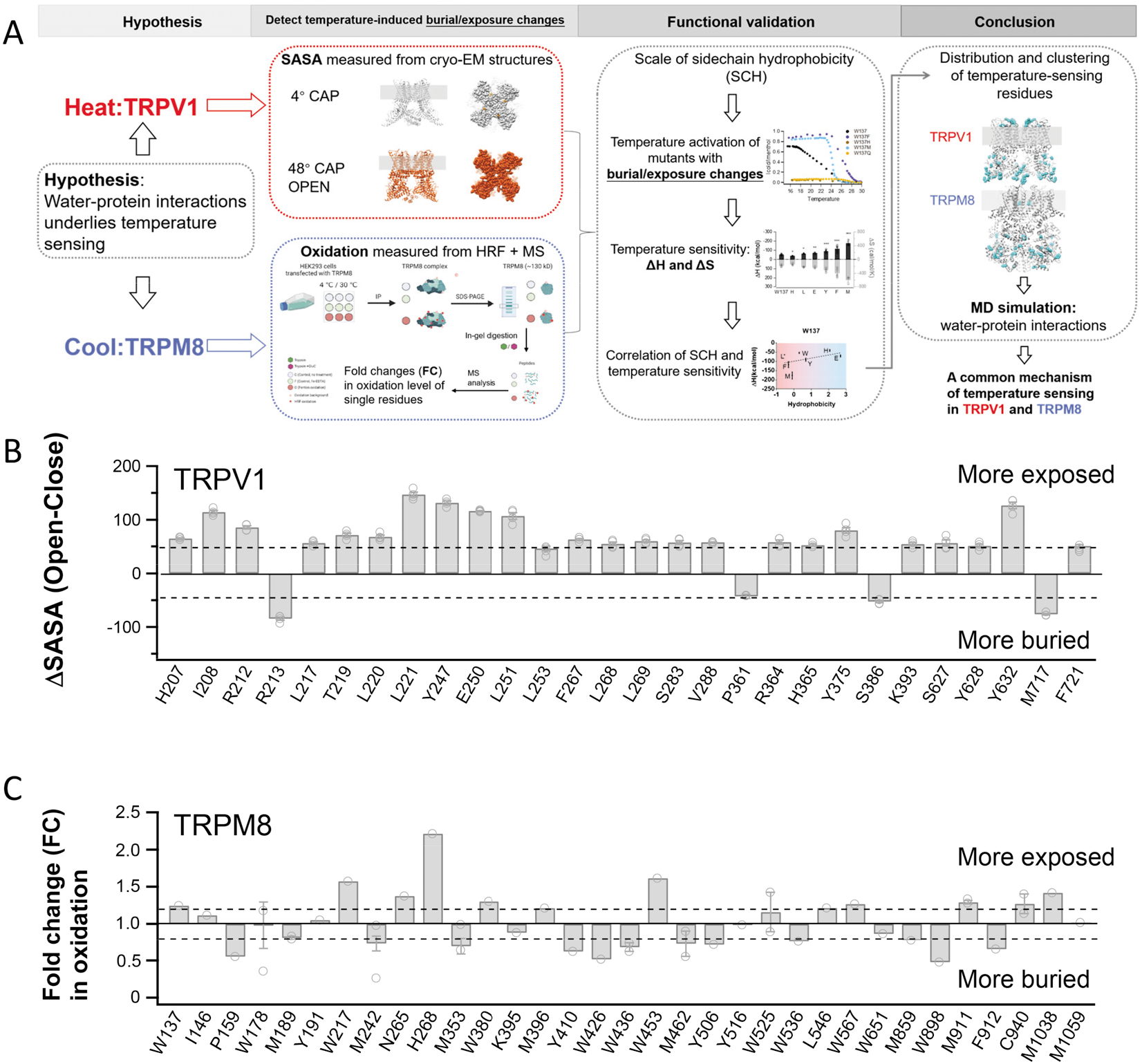
Detecting residues with temperature-induced burial/exposure changes in TRPV1 and TRPM8. (A) The workflow of this hypothesis-driven study, where the hypothesis testing was composed of two major steps: the detecting of temperature-induced burial/exposure changes and the functional validation. (B) From each subunit of high-resolution structure models of TRPV1 channel (open circle in grey) in the closed state at 4℃ (PDB ID: 7LPA) and heat activated state at 42℃ (PDB ID: 7LPE), TRPV1 residues with the changes in SASA measured at least from one subunit being larger than 45 Å^2^ and 50 Å^2^, respectively, as the residues with burial/exposure conformational changes. (C) Bar graph for the FCs showing significant changes in oxidation levels of TRPM8 peptides. FC>1.2 or FC<0.83 under 4℃ and 30℃.FC: fold change. Solid line, 1.0; Dash line, 1.2 and 0.83.

## RESULTS

### Detecting residues with temperature-induced burial/exposure changes

High-resolution structure models of TRPV1 channel in the closed state at 4℃ (PDB ID: 7LPA) and heat activated state at 42℃, though in the presence of capsaicin (PDB ID: 7LPE), have been determined by cryo-EM(Kwon et al., 2021). To identify residues with burial/exposure changes, we directly measured and compared SASA values of each residue in these two models(Fleishman et al., 2011). We further selected residues with the changes in SASA measured at least from one subunit larger than 45 Å^2^ and 50 Å^2^, respectively, as the residues with burial/exposure conformational changes. In this way, we have identified 28 such residues in TRPV1 (Figure 1B, Table 1).

**Table 1.**
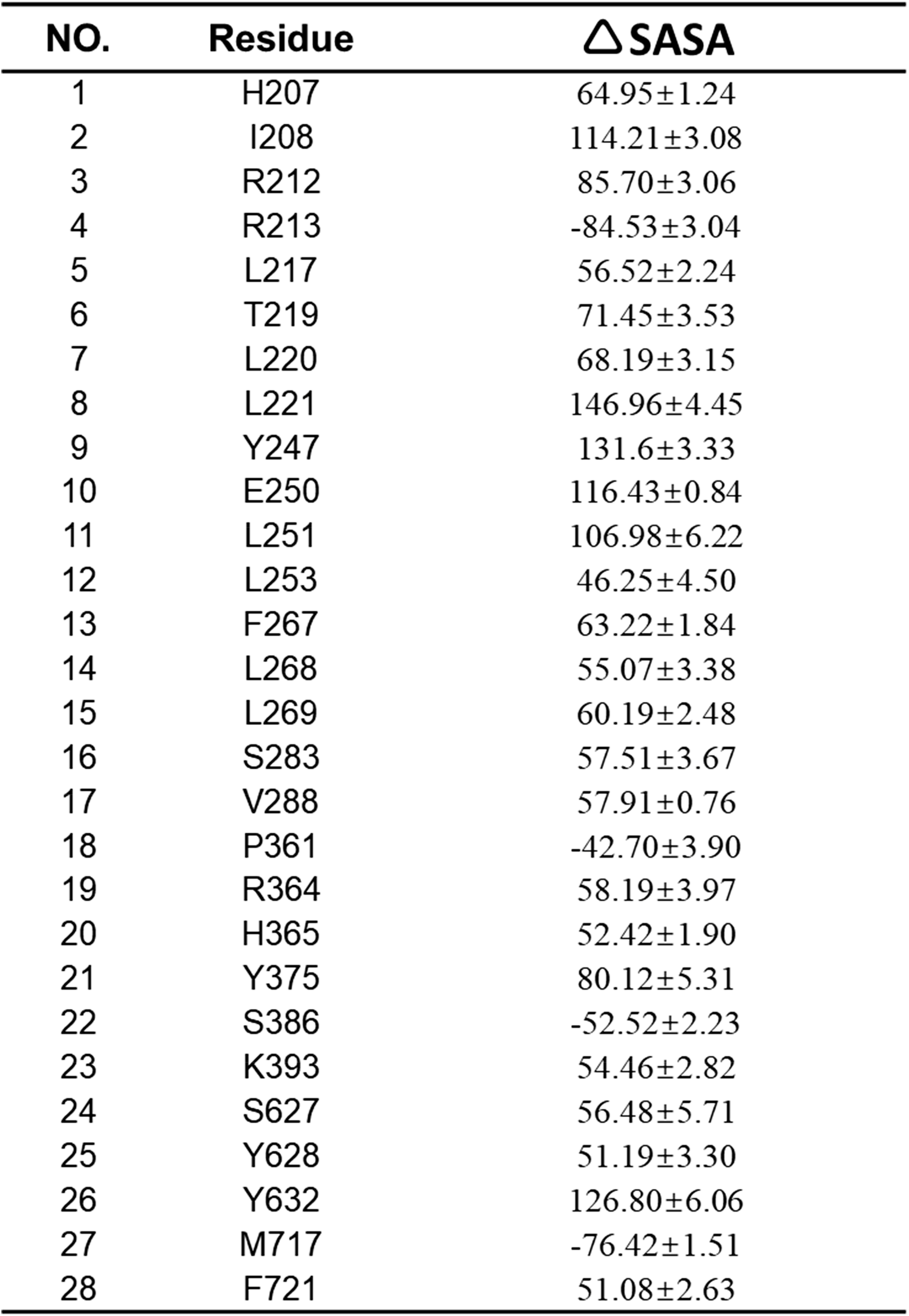
Residues with the changes in SASA larger than 45 Å^2^ and 50 Å^2^, respectively, as the residues with burial/exposure conformational changes. The changes in SASA (ΔSASA) were calculated as the average of four subunits with ± s.e.m.

Unlike TRPV1, though several high-resolution structures of TRPM8 have been resolved by cryo-electron microscopy (cryo-EM) (Diver et al., 2019; Yin et al., 2019; Yin et al., 2018), the cold-activated state of TRPM8 is still lacking. To maximumly identify residues underwent burial/exposure changes during cold activation of TRPM8, we performed an optimized HRF-MS analysis of TRPM8 channel protein expressed in live cells at either 30℃ or 4℃, where TRPM8 remained closed or activated by cold, respectively (Figure 1A, Figure S1 and Methods). In brief, we transiently expressed TRPM8 channel proteins in HEK293 cells. In the living and TRPM8-expressing cells, we generated hydroxyl radicals by an *in situ* Fenton reaction between hydrogen peroxide and Fe(II)-bound EDTA (Zhu et al., 2009a), which can easily diffuse across cell membrane to modify the protein with exposed sidechains. Modern MS technique allowed accurate detection of peptide fragments with a variety of modifications, including oxidation of even a single residue (Aebersold and Mann, 2016). This HRF-MS method has been successfully used to reveal protein higher order structures (Liu et al., 2020) and conformational rearrangements in OmpF channel (Zhu et al., 2009a), Kir channel (Gupta et al., 2010) and EGFR activation (Zhu et al., 2017). Two controls were set as no treatment (Condition C) and treatment with Fe(II)-bound EDTA, respectively. In this way, the sidechains in TRPM8 were oxidized in their native membrane environment before cell lysis, so that the fidelity of information on burial/exposure states in TRPM8 was warranted. In addition, we performed patch-clamp recordings on the TRPM8-expressing cells after Fenton oxidation, where we observed that the channels were still functional (Figure S2).

The analysis of our HRF-MS results was composed of two parts: the TRPM8 spectral library generation and the oxidation quantification. For library generation, TRPM8 and its binding partners were firstly detected by MS acquisition in data-dependent acquisition (DDA) mode and processed into a complex library (Zhu et al., 2020a), which was thereafter transferred as a database to downscale search space. The TRPM8 peptides were acquired in DDA MS mode and identified using the downscaled database (false discovering rate ≤ 0.01) to profile and generate a TRPM8-specific spectral library (Figure S1A). The library covered 83.6% of TRPM8 primary amino acid sequence and contained 1777 peptides and 810 oxidation sites (Figure S1A). An example of the fragmentation spectra of a peptide in the library was presented in Figure S1C.

For quantification of oxidation efficiency, the TRPM8 peptides were acquired by MS in data-independent acquisition (DIA)-MS mode at either 30℃ or 4℃, matched against the spectral library, and quantified according to the summed intensities at MS1 and MS2 (Figure S1B and S1D). By performing MS profiling, we measured the oxidation level of TRPM8 peptides as relative oxidation efficiency (Figure 1C). For instance, we measured the elution peak groups of the matched peptide ^415^AFSTNEQDKDNWNGQLK^431^(Figure S1D), which were used to calculate the relative oxidation efficiency.

We further compared the fold change (FC) in relative oxidation level measured at 4℃ and 30℃. If the FC value of a peptide or residue is larger than 1.20 or smaller than 0.83 (Zhu et al., 2009a; Zhu et al., 2011), we regarded this peptide became significantly more exposed or buried in 4℃, respectively (Figure 1C and Table 2). Only the peptides containing one oxidized residue were considered. In this way, we have identified 33 single residues with cold induced burial/exposure changes from HRF-MS (Table 1, table S1 and table S2).

**Table 2.** Fold change (FC) in relative oxidation efficiency of the residues with buried/exposed changes identified by HRF-MS. FC was calculated as the ratio of relative oxidation efficiency determined at 4℃ to the value measured at 30℃.

| **NO.** | **Residue** | **Fold change (FC) in relative**<br>**oxidation efficiency** |
| --- | --- | --- |
| 1 | W137 | 1.25 |
| 2 | I146 | 1.12 |
| 3 | P159 | 0.55 |
| 4 | W178 | 0.98 |
| 5 | M189 | 0.81 |
| 6 | Y191 | 1.05 |
| 7 | W217 | 1.57 |
| 8 | M242 | 0.73 |
| 9 | N265 | 1.38 |
| 10 | H268 | 2.21 |
| 11 | M353 | 0.70 |
| 12 | W380 | 1.30 |
| 13 | K395 | 0.88 |
| 14 | M396 | 1.21 |
| 15 | Y410 | 0.62 |
| 16 | W426 | 0.52 |
| 17 | W436 | 0.62 |
| 18 | W453 | 1.62 |
| 19 | M462 | 0.72 |
| 20 | Y506 | 0.72 |
| 21 | Y516 | 0.99 |
| 22 | W525 | 1.16 |
| 23 | W536 | 0.76 |
| 24 | L546 | 1.22 |
| 25 | W567 | 1.27 |
| 26 | W651 | 0.87 |
| 27 | M859 | 0.78 |
| 28 | W898 | 0.48 |
| 29 | M911 | 1.29 |
| 30 | F912 | 0.66 |
| 31 | C940 | 1.27 |
| 32 | M1038 | 1.42 |
| 33 | M1059 | 1.02 |

To validate the HRF-MS results, we measured the SASA values from our recently resolved cryo-EM structure of mouse TRPM8 channel in the closed state (a related manuscript (Zhao et al., 2021)) (Figure S3A). The relative oxidation efficiency measured at 30℃, where TRPM8 channel was in the closed state, correlated well with the SASA values (Figure S3B). In contrast, at 4℃ where TRPM8 channel was cold activated into the open state, the relative oxidation efficiency no longer correlated with SASA measured from the closed state structure (Figure S3C). In addition, we incorporated the fluorescent unnatural amino acid ANAP into TRPM8. If the ANAP emission peak shifts to longer wavelength, it indicates the residue becomes more exposed to hydrophilic environment(Chatterjee et al., 2013). Though many sites in TRPM8 did not tolerate ANAP incorporation as we have observed before(Xu et al., 2020; Yang et al., 2020), some ANAP-incorporated mutants remained to be functional and showed shifts in emission spectra during cold activation (Figure S3D to S3F). Residues W436 and M1059 were identified in both HRF-MS and ANAP experiments. Their FC and shifts in ANAP emission peak were in agreement. For instance, at W436 both the FC of 0.62 and left shifts in ANAP emission suggested this site became more buried upon cold activation (Figure S3F). Therefore, both our cryo-EM structure and ANAP imaging of TRPM8 supported the validity of the HRF-MS results.

### Functional validation in TRPV1 mutants

With the identification of residues with burial/exposure changes from cryo-EM structures, we next tested whether altering SCH at these sites could specifically modulate heat sensitivity in TRPV1(table S3). The relative hydrophobicity of amino acids has been measured before (Hessa et al., 2005; Moon and Fleming, 2011). We measured the heat activation of all the mutants by patch-clamp recording (Figure 2). By measuring the temperature dependence of current activation, we constructed Van’t Hoff plots to determine ΔH and ΔS of heat activation process in these mutants as in our previous studies(Yang et al., 2010; Yang et al., 2020). For instance, from SASA measurements (Figure 1B), E250 became more exposed in heat activation. When E250 residue was mutated to increase SCH (E250F, Figure 2A), the ΔH values of the mutants were significantly decreased (Figure 2B). In contrast, from SASA measurements (Figure 1B), P361 became more buried in heat activation. When P361 residue was mutated to increase SCH (P361L, Figure 2D), the ΔH values of the mutants were significantly increased (Figure 2E). We further quantified such a relationship as the slope factor of a linear fitting to the SCH (more negative values shaded in red indicated larger SCH) and ΔH plot (Figure 2C and 2F). Therefore, for a more exposed residue in heat activation like E250, a positive slope factor indicated it supported our water-protein hypothesis on temperature sensing. P361 became more buried in heat activation (Figure 1B), so it exhibited a negative slope factor, which also agreed with our hypothesis on temperature sensing.

**Figure 2.**
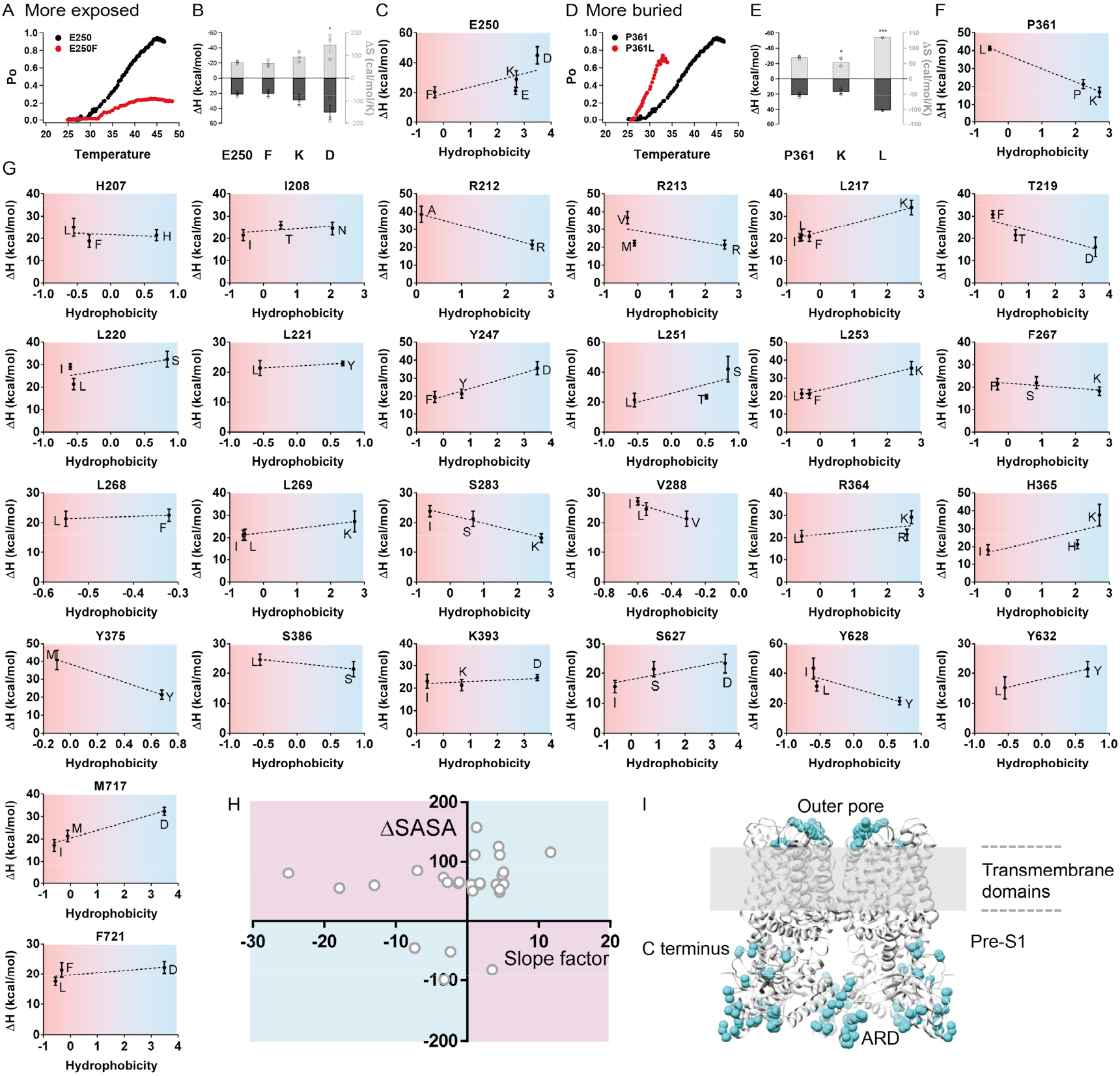
Distribution of TRPV1 residues with correlated changes in SCH and heat sensitivity. (A to C) Representative whole-cell current recordings of heat activation (A), measured ΔH values (filled bars, left axis) and ΔS values (open bars, right axis) (B), correlation between SCH and ΔH values (C) for TRPV1 Y247 residue with more exposed sidechains in heat activation (mean ± SEM; *, ** and *** denote P < 0.05, 0.01 and 0.001, respectively; n = 5-to-7). Residues with large or small SCH (negative or positive Hydrophobicity value) are shaded in red or blue, respectively. (D to F) Representative whole-cell current recordings of heat activation (D), measured ΔH values (filled bars, left axis) and ΔS values (open bars, right axis) (E), correlation between SCH and ΔH values (F) for TRPV1 P361 residue with more buried sidechains in heat activation (mean ± SEM; *, ** and *** denote P < 0.05, 0.01 and 0.001, respectively; n = 5-to-7). (G) Correlation between SCH and ΔH values for TRPV1 residues with changes in SASA larger than 50 Å during heat activation. (H) The slope factor calculated from fitting sidechain hydrophobicity and ΔH values to a linear function for each site with burial/exposure changes was plotted against the ΔSASA value of the corresponding site. The 1^st^ and 3^rd^ quadrant, where the sites followed the predictions from the water-protein interaction hypothesis of cold sensing, were shaded in cyan. The 2^nd^ and 4^th^ quadrant, where the sites did not follow the predictions from the water-protein interaction hypothesis of cold sensing, were shaded in purple. (I) The sites located in the 1^st^ and 3^rd^ quadrant were mapped onto the cryo-EM structure of TRPV1 (PDB ID: 7LPE) with their sidechains shown in cyan.

To reveal the general trend regarding the SCH and heat sensitivity, we performed such analysis on each of the 28 TRPV1 sites with burial/exposure changes identified in cryo-EM structures (Figure 1B and Fig 2G). When we plotted the changes in SASA values (ΔSASA) against the slope factor of these sites, we observed that for a majority of these sites (Figure 2H, 19 out of 28 residues in the 1^st^ and 3^rd^ quadrant shaded in cyan), the burial/exposure conformational changes and SCH correlated well with temperature sensitivity, so that our hypothesis on heat sensing was supported. There were also sites located in the 2^nd^ and 4^th^ quadrant shaded in purple, where changes in heat sensitivity cannot be explained by our hypothesis. When we further mapped these two groups of sites onto TRPV1 channel structure, we observed that the residues supporting our hypothesis were widely distributed in the outer pore, pre-S1, ankyrin-repeat domains and the C terminus (Figure 2I, residues in cyan). Indeed, all regions have been suggested to be critical for heat activation of TRPV1 in previous studies(Grandl et al., 2010; Vlachova et al., 2003; Yang et al., 2010; Yao et al., 2011; Zhang et al., 2018).

### Functional validation in TRPM8 mutants

With the identification of residues in TRPM8 that exhibited burial/exposure changes in cold activation, we also tested whether altering SCH at these sites could specifically modulate cold sensitivity. We mutated each of the residues identified in HRF-MS (Figure 1C and Table 2) to be either more hydrophobic or hydrophilic (table S4).

We then measured the cold activation of all the mutants by patch-clamp recording (Figure 3). For instance, from HRF-MS experiments, W380 became more exposed in cold activation with a FC value of 1.30 (Figure 3A and Table 2). when W380 residue was mutated to increase SCH (W380L, W380I and W380F), the ΔH values of the mutants were largely increased. In contrast, for the W380K and W380D mutants with decreased SCH, their ΔH values were reduced. Therefore, increasing SCH at W380 indeed led to higher cold sensitivity at W380 site. We also quantified such a relationship as the slope factor of a linear fitting to the hydrophobicity and ΔH plot. Therefore, for a more exposed residue in cold activation like W380, a positive slope factor indicated it supported our water-protein hypothesis on temperature sensing. In contrast, residues M242 became more buried in cold activation with a FC value of 0.73 (Figure 3D and Table 2). Its negative slope factor also agreed with our hypothesis on temperature sensing.

**Figure 3.**
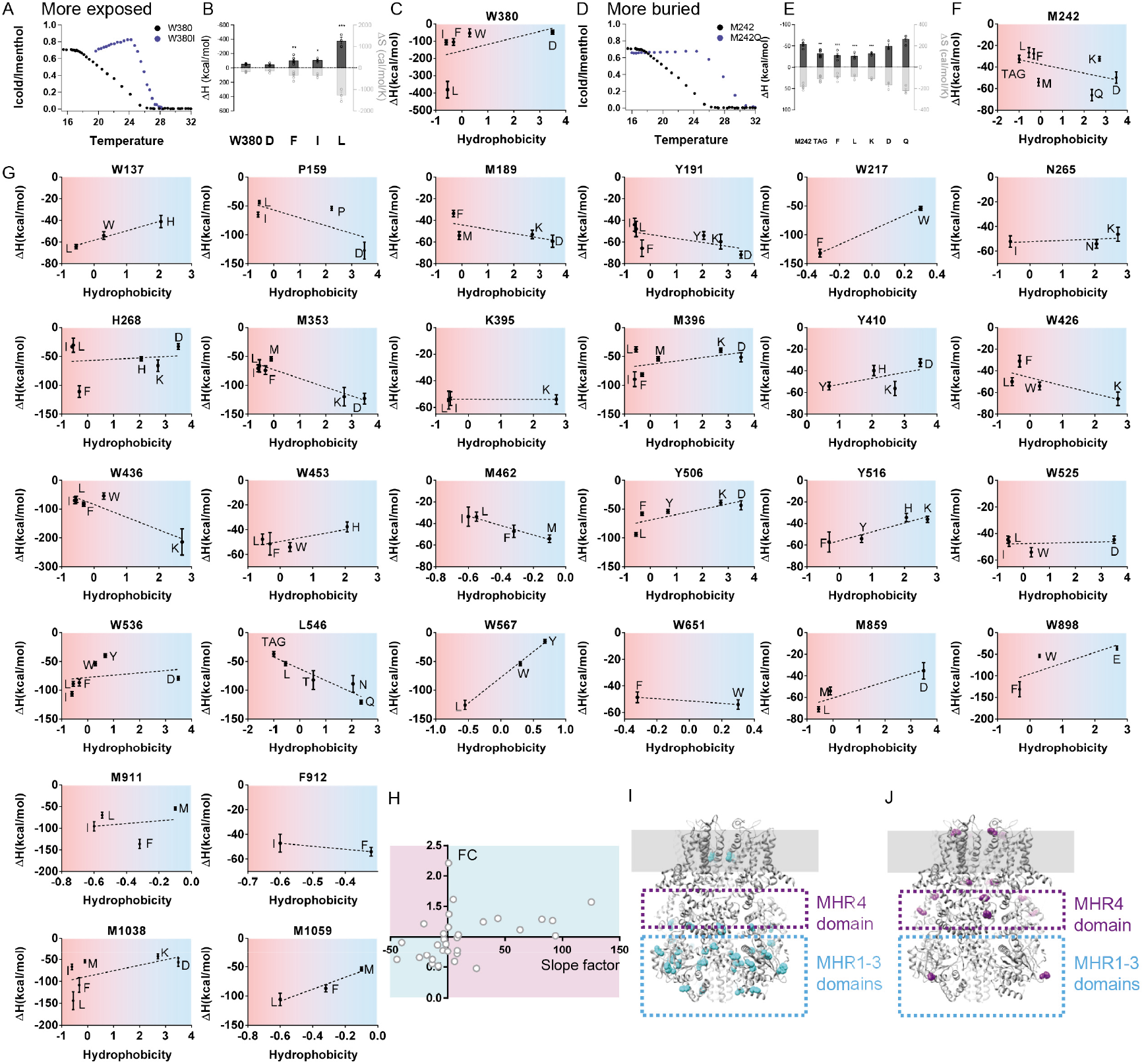
Clustering of TRPM8 residues with correlated changes in SCH and cold sensitivity in the MHR1-3 domains. (A to C) Representative whole-cell current recordings of cold activation (A), measured ΔH values (filled bars, left axis) and ΔS values (open bars, right axis) (B), correlation between sidechain hydrophobicity and ΔH values (C) for TRPM8 W380 residue with more exposed sidechains in cold activation (mean ± SEM; *, ** and *** denote P < 0.05, 0.01 and 0.001, respectively; n = 5-to-7). Residues with large or small SCH (negative or positive Hydrophobicity value) are shaded in red or blue, respectively. (D to F) Representative whole-cell current recordings of cold activation (D), measured ΔH values (filled bars, left axis) and ΔS values (open bars, right axis) (E), correlation between sidechain hydrophobicity and ΔH values (F) for TRPM8 M242 residue with more buried sidechains in cold activation (mean ± SEM; *, ** and *** denote P < 0.05, 0.01 and 0.001, respectively; n = 5-to-6). (G) Correlation between SCH and ΔH values for TRPM8 residues with changes in FC values during cold activation. (H) The slope factor calculated from fitting SCH and ΔH values to a linear function for each site with burial/exposure changes was plotted against the FC value of the corresponding site. The 1^st^ and 3^rd^ quadrant, where the sites followed the predictions from the water-protein interaction hypothesis of cold sensing, were shaded in cyan. The 2^nd^ and 4^th^ quadrant, where the sites did not follow the predictions from the water-protein interaction hypothesis of cold sensing, were shaded in purple. (I and J). The sites located in the 1^st^ and 3^rd^ quadrant, as well as in the 2^nd^ and 4^th^ quadrant, were mapped onto the cryo-EM structure of TRPM8 with their sidechains shown in cyan and purple, respectively.

To reveal the general trend regarding the sidechain hydrophobicity and cold sensitivity, we performed such analysis on each of the sites identified in HRF-MS (Figure 3G). When we plotted the FC values against the slope factor of these sites, we observed that like TRPV1 (Figure 2H), for a majority of these sites (Figure 3H, 21 out of 33 residues in the 1^st^ and 3^rd^ quadrant shaded in cyan), the burial/exposure conformational changes and SCH correlated well with cold sensitivity, so that our hypothesis on temperature sensing was supported. There were also sites located in the 2^nd^ and 4^th^ quadrant shaded in purple, where changes in cold sensitivity cannot be explained by our hypothesis. When we further mapped these two groups of sites onto TRPM8 channel structure, we clearly observed that the residues supporting our hypothesis were clustered within the N terminal MHR1-3 domains (Figure 3I, residues in cyan), while the other residues were located within the MHR4 domain and the other parts of TRPM8 (Figure 3J, residues in purple). Such a distinct pattern in spatial distribution of residues further prompted us to investigate the MHR1-3 domains.

### MHR1-3 domains directly responded to cold

The clustered distribution of residues supporting our hypothesis suggested that the MHR1-3 is critical for cold sensing in TRPM8. If MHR1-3 directly senses cold, these domains should show cold-induced conformational changes. To test this hypothesis, we first expressed and purified the MHR1-3 domains (Figure 4A, residue 109 to 500 in blue) as a soluble protein (Figure 4B). We then monitored the cooling induced conformational changes in MHR1-3 by either far-UV circular dichroism (CD) or intrinsic tryptophan fluorescence measurements, which are classic strategies widely used to study thermodynamic properties of domains in TRP channel (Kim et al., 2020) and other proteins (Greenfield, 2006). The CD spectra of MHR1-3 exhibited two minima at 208 and 222nm (Figure 4C), which was characteristic for α-helix containing proteins like MHR1-3 (Figure 4A). Cooling to 10℃ induced changes in CD spectra, which displayed two-state behavior (Figure 4D). Fitting the data to a two-state sigmoidal model as in a previous study(Kim et al., 2020) yielded a ΔH of −44 ± 2 kcal/mol and a melting temperature of 31.3 ± 0.3 °C (Figure 4D, dots and curve in black). ΔH values measured from TRPM8 open probability in previous studies ranged from −57.8 kcal/mol to −112 kcal/mol (Brauchi et al., 2004; Yang et al., 2010), so it indicates that assuming a full cooperativity, three or less copies of MHR1-3 in the tetrameric TRPM8 channel are sufficient to account for the enthalpic changes in cold activation. Moreover, the melting temperature of MHR1-3 was between the normal body temperature of mammalians like mouse (∼36.5°C) and the cold activation threshold of TRPM8 (∼28°C) (Figure 4D, dots and curve in red), thus enabling the prompt detecting of temperature drops.

**Figure 4.**
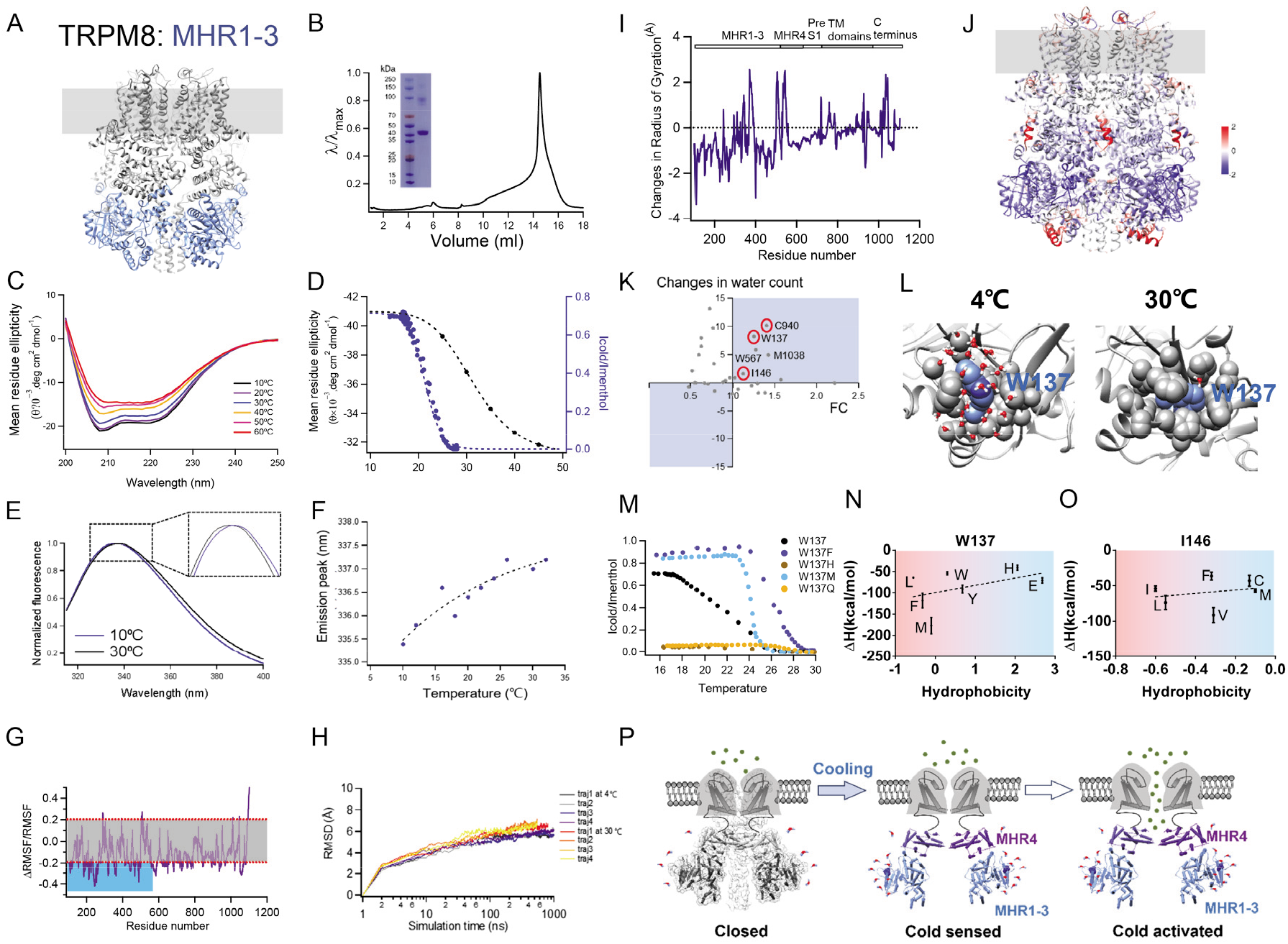
Water-protein interactions for temperature sensing. (A) The location of MHR1-3 domains (colored in blue) in TRPM8. (B) Size-exclusion chromatography of the protein of MHR1-3 on Superose 6 (GE Healthcare) and SDS PAGE. (C) Representative CD spectra of MHR1-3 domains measured at different temperature levels. (D) Comparison of cold activation of TRPM8 current normalized to menthol activation (dots in blue) and changes in mean residue ellipticity (dots in black). (E) Representative intrinsic tryptophan emission spectra of MHR1-3 domains measured at different temperature levels. The inlet showed the cooling induced shifts in emission peak. (F) Temperature dependence of intrinsic tryptophan emission peaks values. (G) RMSD (relative to the TRPM8 structure in the apo state) plots of MD simulation trajectories at either 4℃ or 30℃. All three RMSD curves stabilize within 100 ns. (H) The fractional changes in RMSF (RMSF at 4℃ minus 30℃) normalized to RMSF at 4℃. Dashed lines indicated the 20% significance in fluctuation level, below which any fluctuation was due to nonspecific thermal effects(Wen and Zheng, 2018). (I) The temperature-dependent changes (4℃ minus 30℃) in radius of gyration (Rg) in each residue in TRPM8. (J) The changes in Rg were mapped onto the structure of TRPM8. Negative and positive changes in Rg were colored in blue and red, respectively. The unit of colored scale bar was Å. (K) Changes in contacting water count at residues identified in HRF-MS was plotted against the corresponding FC values. Residues exhibiting consistent changes in both MD and HRF-MS, such as W137 and I146, located in the quadrants shaded in green. (L) Representative snapshots of MD trajectories showing the burial/exposure state of W137 (residue in blue) and the water molecules nearby (oxygen and hydrogen atoms in water colored in red and white, respectively) at 4℃ and 30℃. (M to O) Representative whole-cell current recordings of cold activation (M), correlation between SCH and ΔH values for W136 (N) and I146 (O) residues (mean ± SEM; *, ** and *** denote P < 0.05, 0.01 and 0.001, respectively; n = 4-to-6). Residues with large or small SCH (negative or positive Hydrophobicity value) are shaded in red or blue, respectively. (P) A schematic diagram showing the water-protein interactions as the mechanisms underlying cold sensing in TRPM8.

We further measured the temperature dependence of intrinsic tryptophan emission. There are eight tryptophan residues in MHR1-3, so if temperature induces conformational changes in MHR1-3 to alter the local chemical environment of these residues, their emission peak will shifted (Royer, 2006). We observed cooling from 30°C to 10°C induced blue shifts in intrinsic tryptophan emission (Figure 4E and 4F). In contrast, when the N terminal ankyrin-repeat like domain (ARD) of the heat sensitive TRPV2 channel was expressed and purified (Figure S4A and S4B), within the same 30°C to 10°C temperature range we did not observe temperature dependent changes in either far-UV CD spectra or intrinsic tryptophan emission (Figure S4C to S4F). In addition, when we incorporated ANAP into several sites in MHR1-3, we also observed cooling induced shifts in ANAP emission, suggesting conformational changes occurred there (Figure S3D to S3F). Therefore, our observations suggested that the MHR1-3 domain protein, which hosted the residues changing their burial/exposure state to modulate cold sensitivity in TRPM8 (Figure 3I), directly responded to cold with a sufficiently large enthalpic changes for cold activation. This is also supported by a recent study of TRPM8 evolution, which shows that the formation of the functional MHR1-3 domain in TRPM8 bestowed the channel with cold sensitivity(Lu et al., 2022).

### Energetics in water-protein interactions for temperature activation

To further quantify whether the energetic changes associated with water-protein interactions in thermoTRPs is sufficient for temperature activation, in TRPM8 we directly probe water-protein interactions at residues with burial/exposure changes in the MHR1-3 domains. Due to limitations in resolution, we cannot directly observe water molecules in our cryo-EM structures of mouse TRPM8 (a related manuscript (Zhao et al., 2021)) or other reported TRPM8 structures (Diver et al., 2019; Yin et al., 2019; Yin et al., 2018). Instead, based on our cryo-EM structure in the apo state, we performed multiple all-atom molecular dynamics (MD) simulations at either 4°C or 30°C with a total simulation length of over 6 µs (Figure 4G). Though TRPM8 channel was not directly opened by low temperature during the simulations, we gained unique insights into the cold sensing mechanisms.

MD simulation corroborated our previous findings in the critical role of MHR1-3 in cold activation of TRPM8. Interestingly, we observed that in most domains, the root mean square fluctuation (RMSF) at 4°C was smaller than those at 30°C (Figure 4H, negative ΔRMSF values), indicating that lowering the temperature reduced the mobility of atoms in TRPM8. The MHR1-3 showed the most significant decrease (Figure 4H, shaded area in blue), which was larger than a stringent 20% threshold of non-specific thermal effect (Wen and Zheng, 2018) (Figure 4H, shaded area in grey between solid lines in red). This is also consistent with previous reports that the ΔS is negative in cold activation (Brauchi et al., 2004; Yang et al., 2010). We also measured the changes in radius of gyration (R_g_) between 4°C and 30°C (Figure 4I). The negative changes in R_g_, which suggested the residues became more packed, were also clustered in the MHR1-3 domains (Figure 4I and 4J).

More importantly, we calculated the changes in protein-water electrostatic interaction energy and van der Waals (vdw) energy of the TRPM8 simulation system. We observed a large decrease in the protein-water electrostatic interaction energy (about −84.8 kcal/mol after scaled by a factor of 80, which is the typical value of dielectric coefficient in proteins (Wen and Zheng, 2018)) at 4°C as compared to that at 30°C (Table 4). Interestingly, when the protein-water electrostatic interaction energy in the last 100 ns of simulation was compared to that of the first 100 ns simulation at 4°C, only in MHR1-3 domains we observed a time dependent decrease in protein-water electrostatic energy (−36.9 kcal/mol), while MHR4 and the rest of TRPM8 protein showed an increase (13.8 kcal/mol and 15.6 kcal/mol, respectively). In comparison, at 30°C the change in protein-water electrostatic energy of MHR1-3 was much smaller (−12.3 kcal/mol), indicating such changes were induced by the low temperature. Therefore, our MD results again suggested that it is the MHR1-3, but not MHR4 or other domains in TRPM8, is directly responding to cold.

**Table 3.** Cold and menthol activation of TRPM8 mutants at residue W137, I146 and C940. Temperature was decreased to below 10℃ to activated TRPM8 mutants. A saturating concentration of menthol (2 mM) waw used in ligand activation.

| Mutant | Cold Activation | Menthol Activation |
| --- | --- | --- |
| W137R | NO | NO |
| W137H | NO | YES |
| W137K | NO | NO |
| W137D | NO | NO |
| W137E | YES | YES |
| W137S | NO | NO |
| W137T | NO | NO |
| W137N | Too small to quantify | YES |
| W137Q | Too small to quantify | YES |
| W137C | NO | NO |
| W137G | NO | NO |
| W137P | NO | NO |
| W137A | NO | NO |
| W137V | NO | NO |
| W137I | NO | NO |
| W137L | YES | YES |
| W137M | YES | YES |
| W137F | YES | YES |
| W137Y | YES | YES |
| I146R | NO | NO |
| I146H | NO | NO |
| I146K | NO | NO |
| I146D | NO | NO |
| I146E | NO | NO |
| I146S | NO | NO |
| I146T | NO | NO |
| I146N | NO | NO |
| I146Q | NO | NO |
| I146C | YES | YES |
| I146G | NO | NO |
| I146P | NO | NO |
| I146A | NO | NO |
| I146V | YES | YES |
| I146L | YES | YES |
| I146M | YES | YES |
| I146F | YES | YES |
| I146Y | NO | NO |
| I146W | NO | NO |
| C940K | No | No |
| C940D | No | No |
| C940I | No | No |
| C940L | No | No |
| C940F | No | No |
| C940A | No | No |
| C940Y | No | No |
| C940G | No | No |
| C940S | No | No |
| C940W | No | No |
| C940T | No | No |
| C940R | No | No |
| C940P | No | No |
| C940H | No | No |
| C940N | No | No |
| C940M | No | No |
| C940E | No | No |
| C940V | No | No |
| C940Q | No | No |

**Table 4.** Changes in energy measured from MD simulation of TRPM8. Changes were calculated as energy values at 4℃ minus those at 30℃.

| Interacting components | Electrostatic energy changes (kcal/mol) | Electrostatic energy changes scaled by dielectric coefficient of 80 (kcal/mol) | vdw energy changes (kcal/mol) |
| --- | --- | --- | --- |
| Protein-water interactions in: |  |  |  |
| MHR1-3 | -2432.9 | -30.411 | -146.8 |
| MHR4 | -1725.2 | -21.565 | -125.8 |
| The rest | -2625.0 | -32.812375 | -215.1 |
| Whole TRPM8 protein | -6783.1 | -84.788375 | -487.7 |
| Protein-lipid interactions | 451.6 | 5.64475 | 7.55 |
| Intra-protein interactions | 2233.4 | 27.9175 | -211.6 |

Furthermore, we counted the changes in number of water molecules contacting TRPM8 protein residues by lowering temperature to 4°C (Table 5) and compared the values to FC measured in HRF-MS (Figure 4K). For instance, residue W137 was mostly buried at 30°C (Figure 4L) but cooling to 4°C caused conformational rearrangements, so that W137 became exposed to hydrophilic environment with about eight more water molecules in contact (Figure 4L). FC of W137 measured in HRF-MS was 1.25 (Table 2), again indicating a buried to exposed changes by cooling at this site. Other residues, such as I146 and C940, also showed increase in contacting water (Figure 4K), so that there could easily be more than ten contacting water molecules changed per subunit up cold activation. Assuming hydrogen bond strength ranges from 1.5 kcal/mol to 9.5 kcal/mol (Franks, 2000; Sheu et al., 2003; Yang et al., 2015) and there is only one hydrogen bond formation/breaking per water molecule, a change of ten water molecules may cause an energetic change of 15 kcal/mol to 95 kcal/mol. Therefore, in the tetrameric TRPM8 channel with four copies of burial/exposure residues, the estimated energy changes associated with changes in water count (48 kcal/mol to 304 kcal/mol) is sufficient for the ΔH as measured from TRPM8 open probability in previous studies (Brauchi et al., 2004; Yang et al., 2010).

**Table 5.** Changes in water counts measured from MD simulation of TRPM8. Changes were calculated as water counts at 4℃ minus those at 30℃. Water molecules within 4.5 Å of the sidechain in a residue were counted.

| NO. | Residue | Change in water counts (4°C – 30°C ) |
| --- | --- | --- |
| 1 | W137 | 8.26 |
| 2 | I146 | 1.66 |
| 3 | P159 | 0.05 |
| 4 | W178 | 4.97 |
| 5 | M189 | 0.96 |
| 6 | Y191 | 6.04 |
| 7 | W217 | 0.56 |
| 8 | M242 | 3.41 |
| 9 | N265 | 2.57 |
| 10 | H268 | -1.74 |
| 11 | M353 | 0.00 |
| 12 | W380 | -0.17 |
| 13 | K395 | -1.76 |
| 14 | M396 | 0.15 |
| 15 | Y410 | 5.00 |
| 16 | W426 | 7.99 |
| 17 | W436 | 2.18 |
| 18 | W453 | 6.50 |
| 19 | M462 | 0.40 |
| 20 | Y506 | 13.18 |
| 21 | Y516 | -1.93 |
| 22 | W525 | -4.56 |
| 23 | W536 | 9.74 |
| 24 | L546 | -0.44 |
| 25 | W567 | 5.90 |
| 26 | W651 | 1.67 |
| 27 | M859 | -1.42 |
| 28 | W898 | -0.68 |
| 29 | M911 | -0.09 |
| 30 | F912 | 0.90 |
| 31 | C940 | 10.16 |
| 32 | M1038 | 4.97 |
| 33 | M1059 | 0.93 |

To further validate our hypothesis, we mutated W137, I146 and C940, in which both MD and HRF-MS suggested an increase in water contact (Figure 4K, circles in red), to all the other 19 residues and characterized each of the mutants by patch-clamp recording. We found that none of the 19 mutants at site C940 was function, while there were only eight and five mutants at the sites 137 and 146 remained to be functional, respectively (Table 3). The cold activated current of W137N and W137Q was too small to accurately measure, so that they were excluded from analysis. Like our previous observation (Figure 3), most of the mutants followed the prediction from our hypothesis: increasing SCH at these exposed sites caused enhancements in cold sensitivity (Figure 4M to 4O). Therefore, based on all the observations above, we believe that our proposed water-protein interactions as mechanisms of cold sensing in TRPM8 is supported (Figure 4O).

In TRPV1, we previously have simulated its heat activation process without adding capsaicin in all-atom molecular dynamics (Wen and Zheng, 2018). By comparing the closed state of TRPV1 determined by cryo-EM and the heat-driven open state predicted in our MD simulation, we observed a large increase in the protein-water electrostatic interaction energy (74.7 ± 6.4 kcal/mol after scaled by the factor of 80 for dielectric coefficient in proteins), which is with the range of experimentally measured ΔH in TRPV1 heat activation (26.8 to 100 kcal/mol) (Yang et al., 2010; Yao et al., 2010). Therefore, in both TRPM8 and TRPV1, energetic changes in water-protein interactions are sufficient for temperature-driven activation.

## DISCUSSIONS

In this study, we tested our hypothesis of water-protein interactions underlying the temperature sensing in TRPV1 and TRPM8 by exhaustively searching for residues with burial/exposure changes in temperature activation (Figure 1). We systematically altered the SCH in all the identified residues, and then measured the impact of SCH on temperature sensitivity in mutants by patch-clamp recording (Figure 2 and Figure 3). As for TRPM8, these experiments led us to identify the MHR1-3 domains that directly responded to cold, which exhibited clear changes in interactions with water molecules and sufficiently large energetic changes for cold activation. Therefore, we believe that the MHR1-3 domains, especially the residues with burial/exposure changes in these domains, serve as the driving “engine” for cold sensing in TRPM8.

Nevertheless, the residues supporting our hypothesis but distributed outside the intracellular MHR1-3 domains (like M911 in the pore region, Figure 1C and Figure 3G) indicated that a more complex nature of cold sensing and activation mechanism. Indeed, by analyzing TRPM8 channels in various vertebrates, previous studies showed that residues in the transmembrane domains can largely modulate cold activation properties(Matos-Cruz et al., 2017; Pertusa et al., 2018; Yang et al., 2020). However, these analyses were limited to evolutionarily advanced vertebrate species, where cold activation of TRPM8 was modulated but not abolished. As MHR1-3 serves as the engine for cold sensing in TRP8, these domains may also be critical for molecular evolution. Indeed, a recent study shows that during the evolution of terrestrial tetrapods, the formation of the functional MHR1-3 domain in TRPM8 bestowed the channel with cold sensitivity(Lu et al., 2022). Therefore, we believe that these residues located outside the MHR1-3 work as modulators for cold sensing and activation in TRPM8.

In TRPV1, we observed that at the residues with a decrease in SASA during heat activation, by increasing the SCH the temperature sensitivity of the mutant was increased (Figure 2), indicating like TRPM8, a common mechanism of water-protein interactions underlying temperature sensing is also applicable to TRPV1. However, unlike the clustering of residues involved in temperature sensing in the MHR1-3 domains in TRPM8, these residues in TRPV1 were widely distributed in the outer pore, pre-S1, ankyrin-repeat domains and the C terminus (Figure 2I). All these regions have been reported to be important for heat activation of TRPV1 (Grandl et al., 2010; Vlachova et al., 2003; Yang et al., 2010; Yao et al., 2011; Zhang et al., 2018). Our observations in TRPV1 agreed with the theoretical analysis suggesting that the temperature sensing apparatus in TRPV1 could be distributed on a single-residue basis (Clapham and Miller, 2011).

Heat-induced global conformational changes have been revealed in the cryo-EM studies of TRPV1 and TRPV3(Kwon et al., 2021; Nadezhdin et al., 2021), where the heat activation was initiated from the intracellular N and C terminal domains and conformational changes in the outer pore also occurred. To fully establish our proposed mechanism of temperature sensing in thermoTRPs, resolving more channels, including TRPM8, in a temperature-driven open state are needed.

Elucidating the temperature sensing mechanisms in thermoTRPs is crucial for translational applications. thermoTRPs are known to be important pain detectors so that they are targets for analgesic drugs (Julius, 2013). However, many blockers of these channels developed for analgesia caused changes in body temperature and blunt of acute temperature sensation in patients, leading to failures in clinical trials (Gavva, 2009; Kort and Kym, 2012). Understanding the temperature sensing mechanisms and temperature-induced conformational changes will help develop modality-specific blockers, like the cyclic peptide DeC-1.2 we recently designed to inhibit the ligand gating of TRPM8 without affecting cold activation (Aierken et al., 2021), to minimize adverse side effects while exerting analgesic effects.

## ACKNOWDGEMENTS

We are grateful to our lab members for assistance and discussion. We would like to thank Dr. Cheng Ma (Protein Facility of the Core Facility, Zhejiang University School of Medicine) for constructive advice on protein purification. This study was supported by National Natural Science Foundation of China (32122040, 31971040 and 31800990 to F. Y.; 81972492 and 21904107 to T.G.), Zhejiang Provincial Natural Science Foundation of China (LR20C050002 to F.Y.; LR19C050001 to T.G.), and Hangzhou Agriculture and Society Advancement Program (20190101A04 to T.G.). This work was supported by Alibaba Cloud.

## AUTHOR CONTRIBUTIONS

L.X., X.L., W.Z., Z.X., F.Z., X.Y., L.H. and X.C. conducted the experiments including protein expression and purification, HRF-MS and patch-clamp recordings; B.L., B.Z., Y.W., Z.D., and H.W. performed computational studies; F.Y., Y.Z., T.G. and H.W. conceived and supervised the project and prepared the manuscript; F.Y., Y.Z., T.G. and H.W. participated in data analysis and manuscript writing.

## DATA AVAILABILITY

All data needed to evaluate the conclusions in the paper are present in the paper and/or the Supplementary Materials. The proteomic data are deposited in ProteomeXchange Consortium.

(https://www.iprox.cn/page/PSV023.html;?url=1634831300401jcmF with the password: UhEf). Additional data available from authors upon request.

## Supplemental Information

### EXPERIMENTAL PROCEDURES

#### Cell treatment and Fenton oxidation *in vivo*

The transfected HEK cells were stimulated in three different conditions for 5min: i), C for Ctrl, HEK cells were bathed in 5mL of 1x PBS, no other treatment; ii), F for Fe treatment, namely HEK cells were bathed with 10 mM Fe(II)-bound EDTA in 5mL 1x PBS; and iii), O for OH radical oxidation of cells, which means HEK cells were oxidized by Fenton oxidation *in vivo*. The cell density of these above conditions was around 10^7^ cells/ml as optimized previously (*23*). The oxidation process was slightly modified from OMPF footprinting as published previously (Zhu et al., 2009b).

Firstly, HEK cells were resuspended and bathed in 5mL 1x PBS. Then 10mM (NH4)_2_Fe(SO_4_)_2_ and 25mM Na_2_EDTA were added into the system and mixed by brief votexing, then 70 μL of 30% H_2_O_2_ were added to make a final concentration of 0.3% H_2_O_2_ into the suspension for Fenton oxidation, lasting for 5min. Two temperatures were included in this study for oxidation under ion channel opening (4℃) and ion channel closure (30℃). The reaction was quenched after adding equal volume of Tris-HCl (100mM, pH 8.0, Sangon Biotech, B548127). The cells were collected by centrifugation at 1100 rpm, room temperature for 10 min for later processes. For each condition, three biological replicates were conducted. As a summary, HEK cells treated under three biological conditions at two temperatures were included in this study.

#### Immunoprecipitation (IP), SDS PAGE separation, and in-gel digestion

The detailed protocols for IP, SDS PAGE separation, and in-gel digestion were specified as previously published (Zhu et al., 2011; Zhu et al., 2017). Briefly, the treated HEK cells were firstly lysed in the in-house prepared lysis buffer (1% Triton X-100, 1% deoxycholate, 0.1% SDS, 150mM NaCl in 50mM Tris-HCl, pH7.4). Thereafter, Streptactin Bead 4FF antibody (SA053005, Smart-Lifesciences) was added to the supernatant over night at 4℃. The bound complexes were pulled down and were subsequently subjected to electrophoresis on an 8% SDS-PAGE. After SDS-PAGE separation, the page was stained with Coomassie blue (A100615-0005, Sangon Biotech, China) to locate the position of TRPM8 protein.

For the identification of TRPM8 and its binding partners, the whole lane was cut into 10 fractions for later in-gel digestion steps. For identification and quantification of TRPM8 only in the latter experiment, only the area around 130 kDa from the lane was selected and cut into 2-3 gel bands. They were subsequently digested into peptides either by i), a two-time trypsin digestion (HLS TRY001C, Hualishi Technology, enzyme: protein ratio at 1:100 each time, 4h plus 12h intervals) at 37℃ overnight; or ii), a sequential two-enzyme digestion first with trypsin (enzyme: protein ratio, 1:100) at 37℃ for 4h, and second with Glu-C (HLS GLU001, Hualishi Technology, (enzyme: protein ratio, 1:100)) at 37℃ overnight. The peptides were extracted, and desalted using C18 spin tips (Pierce^TM^ 84850, Thermo Scientific) according to the manufacturer’s protocol.

#### LC-MS/MS analysis

The LC−MS/MS analysis was performed either on a QE-HF or QE HF-X mass spectrometer (Thermo Fischer) with the same MS setting (table S1) coupled with an UltiMate 3000 liquid chromatography system (Thermo Fischer). Around 500 ng of peptides prepared from each fraction were separated in a fused silica column (15 cm × 75 μm ID, National Institute of Biological Sciences) packed with C18 (1.9 μm 100 Å) at a flow rate of 300 nL/min. Buffer A (2% ACN, 0.1% FA) and buffer B (80% ACN, 0.1% FA) were used for the LC gradient. Nano spray (NSI) was applied for ionization.

For library generation, firstly, DDA MS data acquisition for the 10 fractions of TRPM8 complex was performed at an effective 60 min LC gradient that ramped from 3% to 28% buffer B. This is to identify TRPM8 complex proteins. Secondly, DDA MS data acquisition for the 2-3 fractions around 130kD area of both oxidized and non-oxidized TRPM8 proteins was performed at a longer LC gradient, namely, an effective 90min LC gradient that ramped from 3% to 28% buffer B in 90 min (Zhu et al., 2020b). This is to better characterize TRPM8 oxidation sites. A shotgun DDA-MS strategy was applied with the following parameters: The survey scan (400-1200 m/z) was acquired at a resolution of 60k, a maximum ion accumulation time of 80 ms, and an AGC target of 3e6. The top 20 ions were selected for MS2 analysis at a resolution of 30k, a maximum ion accumulation time of 100 ms, and an AGC target of 1e5. Dynamic exclusion was activated with a duration of 30 s. Ions with charge state of +1 and >5 were excluded from MS/MS. The isolation width was set to 1.6 Da and the normalized collision energy (NCE) was set at 27. Spectra were acquired in centroid format.

For proteomic identification and quantification of oxidized domains, TRPM8 peptides were spiked in with 10% iRT peptides (Biognosys) for latter calibration. DIA MS data acquisition was performed at an effective 60 min LC gradient that ramped from 3% to 28% buffer B. A DIA-MS strategy was applied (Zhu et al., 2020b). The survey scan (390-1010 m/z) was acquired at a resolution of 60k, a maximum ion accumulation time of 80 ms, and an AGC target of 3e6. The MS2 scan was acquired in 24 windows set as 389.5-410.5, 409.5-430.5, 429.5-450.5, …-749.5-770.5, 769.5-830.5, 829.5-890.5, 889.5-950.5, 949.5-1010.5. The resolution for MS2 was set as 1e6, with an automated maximum ion accumulation time, and an AGC target of 1e6. The normalized collision energy (NCE) was set at 28. Spectra were acquired in centroid format.

#### TRPM8 spectral library generation

In total, 45 DDA files were acquired for library generation, including 28 using trypsin and 17 using trypsin coupled with Glu-C as digestion enzymes, respectively. We also included 28 test samples of TRPM8 under different oxidation conditions during the experiment to increase sequence coverage, as labeled in table S1.

The 10 DDA files of the TRPM8 complex were searched against the fasta sequence composed of the human reviewed fasta, the Pirt sequence, and the *Mus musculus* TRPM8 sequence, using pFind software with the deafult settings and an FDR cutoff of 0.01. Proteins that do not interact with TRPM8 (KEGG: mmu04750) and fulfill one of the following criteria were excluded from the resultant matrix: 1) with only one identified peptide; 2) with multiple matches (protein groups); 3) keratin. The filtered result was transferred to a filtered fasta database.

The 35 DDA files of TRPM8 under different treatment conditions were searched against the filtered fasta database, using pFind software with Open Search settings and an FDR cutoff of 0.01. The following oxidation modifications were considered as variable modifications with maximal modification number per peptide as three:16Ox[M];16Ox[D];16Ox[F];16Ox[H];16Ox[K];16Ox[N];16Ox[P];16Ox[R];1 6Ox[W];16Ox[Y];32Ox[C];32Ox[F];32Ox[M];32Ox[W];32Ox[Y];14Ox[I];14Ox[E];14Ox[K];14Ox[L];14Ox[P]. Other parameters were set as default. Proteins with modifications other than oxidation, carbamidomethyl, or deamination were excluded from the resultant matrix. The filtered result contains 1777 peptide precursors including 810 oxidized sites and was used as the TRPM8 library.

#### DIA MS quantification of oxidized peptides of TRPM8

Total 72 DIA files were used for the DIA quantification, including 36 using trypsin and 36 files using trypsin coupled with Glu-C as digestion enzymes, respectively. The DIA files of TRPM8 were analyzed using Skyline (version: daily-20.2.1.215). The TRPM8 library was imported for spectra match. iRT were imported to calibrate retention time and MS error PPM was set to be lower than 15 ppm. Other parameters were set as default. All fragments were selected for quantification of its origin peptide precursor. For each peptide, the raw intensity at both MS1 and MS2 were added together. A resultant peptide matrix was generated as shown in table S1.

The oxidation level was calculated by:

*Oxi% =Intensity_Oxi_ /(Intensity_Oxi_+ Intensity_nake_)*

The relative oxidation level was calculated by:

Relative Oxi% =Oxi%*_oxi_* /(Oxi%*_ctrl1_* + Oxi%*_ctrl2_*)

The fold changes were calculated by:

*Fold change = Relative Oxi%_4℃_ /Relative Oxi%_30℃_*

#### MS raw data deposition

The proteomic data are deposited in ProteomeXchange Consortium. (https://www.iprox.cn/page/PSV023.html;?url=1634831300401jcmF with the password: UhEf).

#### SASA Calculation

SASA of each residue in TRPV1 structures models was measured by RosettaScripts within the Rosetta suite(Fleishman et al., 2011). SasaMetric SimpleMetric was performed within the Rosetta suite. The scripts to perform SASA measurements were attached in Supplementary Methods.

All the molecular graphics of TRPV1 and TRPM8 models were rendered by UCSF Chimera software version 1.12.

#### Gene synthesis and mutagenesis

*Mus musculus trpm8* (GeneID: 171382) was synthesized by Tsingke (Beijing, China) based on the predicted gene sequence and subcloned into the pEGFP-N1 vector. Murine TRPV1 (gifts from Dr. Michael X. Zhu at University of Texas Health Science Center at Houston) was used in this study. Point mutations were constructed using 2×Taq Plus Master Mix (Vazyme, China) following the manufacturer’s instruction. All point mutations were summarized in table S3 and table S4. All point mutations were confirmed by sequencing.

#### Cell transient transfection

HEK293T cells were cultured in Dulbecco’s modified eagle medium (DMEM) supplemented with 20 mM L-glutamine and 10% fetal bovine serum, incubated at 37°C with 5% CO_2_. Cells were transiently transfected by Lipofectamine 3000 (Life technologies) following manufacturer’s protocol. Patch-clamp recordings were performed 24h after transfection.

#### Electrophysiology

Patch-clamp recordings were performed with a HEKA EPC10 amplifier controlled by PatchMaster softerware (HEKA). Whole-cell recordings were performed at ± 80 mV. Patch pipettes were prepared from borosilicate glass and fire-polished to resistance of ∼4 MΩ. For whole-cell recording, serial resistance was compensated by 60%. A solution with 130 mM NaCl, 10 mM glucose, 0.2 mM EDTA and 3 mM Hepes (pH 7.2) was used in both bath and pipette for whole-cell recordings. Current was sampled at 10 kHz and filtered at 2.9 kHz.

Gravity-driven system (RSC-200, Bio-Logic) was used to perfuse bath solution or menthol onto the cell membrane. Bath and ligand solution were delivered through separate tubes to minimize the mixing of solutions. Patch pipette was placed in front of the perfusion tube outlet.

#### Temperature control

We used a precooled bath solution to activate TRPM8. Cells were recorded and placed in bath solution over 30°C before recording. We used a preheated bath solution to activate TRPV1. Hot bathing solution were maintained at expected temperature with an SH-27B in-line solution heater controlled by a TC-324C temperature controller (Warner). A TA-29 miniature bead thermistor (Harvard Apparatus) was placed right next to the pipette to ensure accurate monitoring of local temperature.

#### Calculation of ΔH and ΔS

To calculate the change of enthalpic (ΔH) and the change of entropic (ΔS) due to the temperature-driven transition, we constructed Van’t Hoff plots and fitted them with the equation:

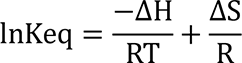

where R represents the gas constant, T represents the temperature in Kelvin, Keq represents the equilibrium constant.

#### Protein expression and purification

Dialysis-based cell free protein synthesis reactions were performed using modified *E.coli* cell extracts, amino acid mix and reaction mix etc. as per described in manufacturer’s (GZL Bioscience Co. Ltd) protocol. PCR product containing target gene and protein expression components (including 5’ T7 promoter, ribosome binding site, start codon and 3’T7 terminator) was added into cell free synthesis reaction as template. The reactions were carried out at 30 °C for 12 hours. Samples were analyzed by SDS-PAGE.

#### Far-ultraviolet circular dichroism (Far-UV CD)

The Far-ultraviolet circular dichroism measurements were performed on 0.2 mg/ml samples in 20 mM Hepes (pH 8.0), 150 mM NaCl and 10% glycerol. The CD spectra were obtained by a circular dichroism spectrometer (Chirascan V100) with a path length cell of 1.0 mm. All experiments were measured from the range of 200 nm to 250 nm, and scanned with an increase of 2°C in the range of 10°C to 50°C. Convert The unit of the CD value is converted into the mean residue ellipticity with the equation:

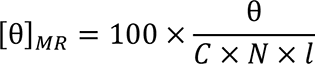

where [θ]_MR_ is the mean residue ellipticity, θ is the ellipticity in millidegrees, *C* is he concentration of protein in molarity (M), *N* is the number of amino acid residues of protein, and *l* is the path length in centimeters.

#### Temperature-dependent intrinsic tryptophan fluorescence

Fluorescence emission spectra were measured on a Spectrofluorometer using 295 nm excitation to reduce the contribution of tyrosine residues. All the samples were prepared in a 0.2 mg/ml solution containing 20 mM Hepes (pH 8.0), 150 mM NaCl and 10% glycerol. The temperature was controlled over a range of 10 °C-30°C in 2 °C increase by using a water circulation system.

#### Preparation of mouse TRPM8 Cryo-EM Structure for MD simulation

To complete missing parts in the cryo-EM structure of mouse TRPM8 in the apo state, this initial model was reconstructed based on SWISS-MODEL (Waterhouse et al., 2018). Four phosphatidylinositol lipids were added to previously proposed lipid-binding sites (Yin et al., 2019). The initial model was fitted into the cryo-EM density map using the MDFF method (Trabuco et al., 2008). CHARMM36m force field (Huang et al., 2017) was used. The initial model was first optimized with a 1000-step energy minimization using the conjugated gradient with line-search algorithm. Then the system was simulated for 10 picoseconds. The simulation temperature was maintained at 300 K using the Langevin algorithm(Zwanzig, 1973) with a damping coefficient of 1 ps^-1^. A scaling coefficient of 0.4 was used to reduce the forces derived from the MDFF grid potential. Chirality restraints, cis-peptide restraints, and secondary structure restraints were applied to the protein during MDFF fitting. Generalized-Born implicit solvent model (Ghosh et al., 1998; Tsui and Case, 2000) was used to describe the solvation effects. A time step size of 0.2 fs was used. For nonbonded interactions, a cutoff distance of 16 Å and a switching distance of 15 Å was used to reduce the computational cost.

#### MD simulation setup

With the MDFF refined model, we used the Membrane Builder function (Jo et al., 2007) of the CHARMM-GUI webserver (Jo et al., 2008; Lee et al., 2016) to embed TRPM8 model in a 180Å×180Å bilayer of 1-palmitoyl-2-oleoyl phosphatidylcholine (POPC) lipids, the lower leaf was mixed with 10% PI(4,5)P_2_, surrounded by a box of water and ions with a buffer distance of 15 Å. To ensure 0.15 M ionic concentration and zero net charge, 514 Na^+^ and 322 Cl^-^ions were added. The aforementioned 4 PI(4,5)P_2_ molecules were kept. OpenMM7.5.1 (Eastman et al., 2017) and CHARMM36m force field (Huang et al., 2017) was used to perform MD simulation on single V100 graphics processing unit (GPU). After energy minimization, six steps of equilibration were performed (with gradually reduced harmonic restraints applied to protein, lipids, water, and ions). Finally, we conducted production MD runs in the NPT ensemble for 1000ns and 500ns at 4°C and 30°C, respectively, four replica were performed for each temperature. MonteCarloMembraneBarostat in OpenMM was applied at 1 bar with an update frequency of every 100 steps. The particle mesh Ewald method (Darden et al., 1993) was used for electrostatics calculations. Analysis and visualization was performed using VMD (Humphrey et al., 1996).

#### RMSF analysis

To assess the flexibility of TRPM8 at individual residue positions during our MD simulation, we calculated the root mean square fluctuation (RMSF) as follows: first, we saved 1600 snapshots from first 500-ns MD trajectories of all low and high temperature simulation respectively (with the first 100 ns of each trajectory discarded) to build the low temperature and the high temperature ensembles; second, we superimposed the Cα coordinates onto the initial structure with a minimal root mean square deviation (RMSD); finally, we calculated the following RMSF at residue position *n* within the range: 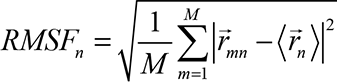, where 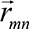 is the Cα position of residue *n* in snapshot *m*, 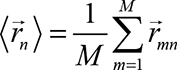 is the average Cα position of residue *n*, and *M* is the total number of snapshots in each ensemble. We then calculated the average of *RMSF_n_* for four equivalent residue positions *n* of the tetramer as the final RMSF for each residue.

#### R_g_ analysis of inward/outward motions

To probe inward/outward motions in TRPM8 at the residue level of details, we used the *measure* command of the VMD program (Humphrey et al., 1996) to calculate the radius of gyration (R_g_) based on the Cα atoms. Since the tetrameric channel was aligned with respect to the Z-axis, the R_g_ essentially describes the average distance between each site to the center of the channel in X-Y plane. Based on the equilibrium MD simulations of the low and high temperature (between 100 and 500 ns), we averaged R_g_ over four MD trajectories, and then computed the change of average R_g_ (n) from T=30°C to 4°C to quantify the cold-activated expansion/contraction at residue position n.

#### Energetic analysis of non-bonded interaction energy

We used the NAMD (Phillips et al., 2005) Energy plugin of the VMD program (Humphrey et al., 1996) to calculate nonbonded energy [including van der Waals (vdW) and electrostatic energy] in the TRPM8-water-membrane system. A 10-Å switching distance and a 12-Å cutoff distance were used for the nonbonded interactions. The CHARMM36m force field was used for the nonbonded parameters.

#### Statistics

All experiments have been independently repeated for at least three times. All statistical data are given as mean ± s.e.m.. Two-sided Student’s *t*-test was applied to examine the statistical significance. N.S. indicates no significance. *, ** and *** indicate *p* < 0.05, *p* < 0.01 and *p* < 0.001, respectively.

### Additional Supplemental Figure

**Figure S1.**
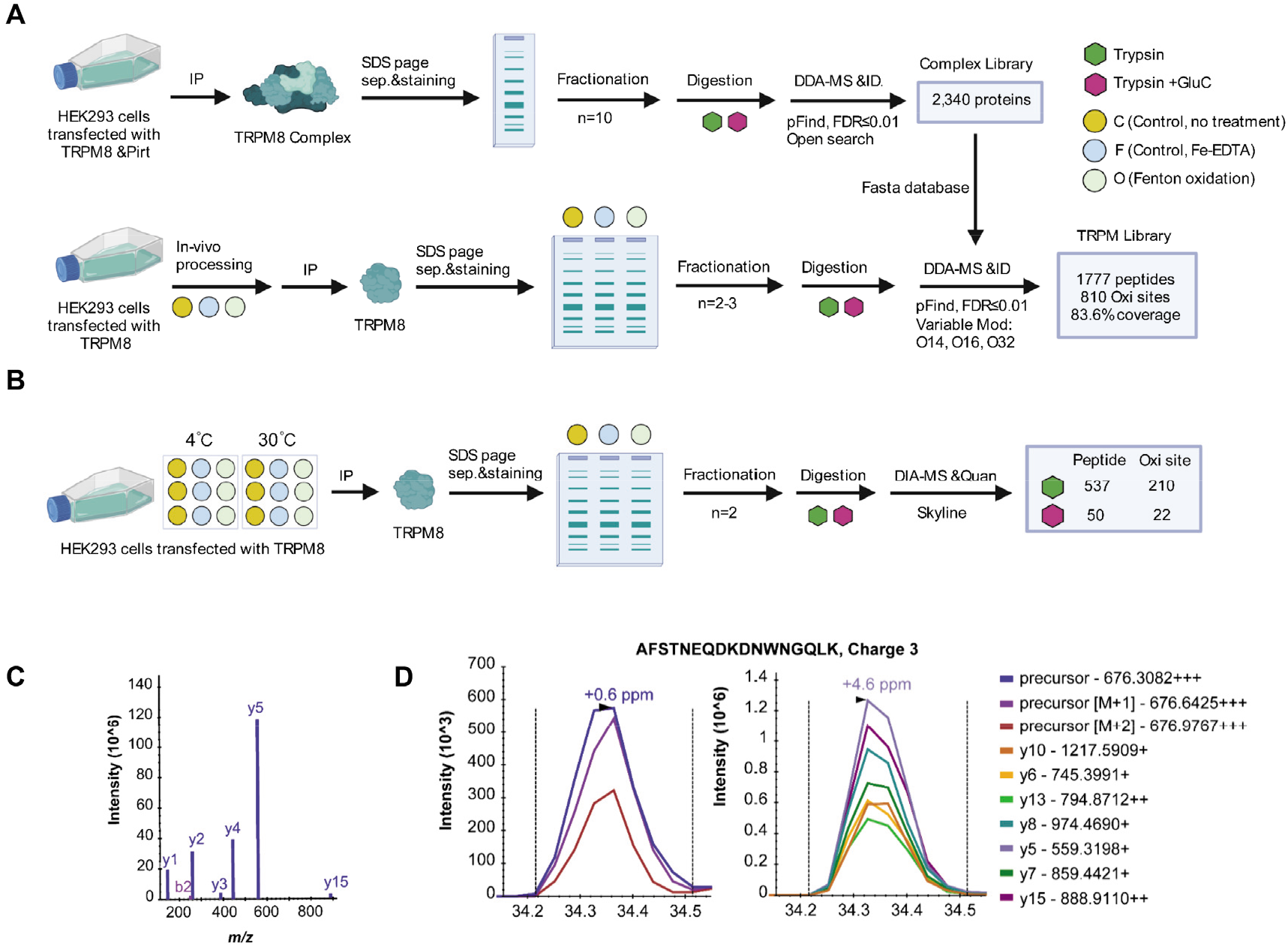
HFR-MS for residues with solvent accessibility changes in TRPM8. (A) The workflow for TRPM8 spectral library generation. IP: immune-precipitation; DDA: data-dependent acquisition; O14: +13.979265; O16: +15.994915; O32: +31.989828. (B) The workflow for TRPM8 oxidation site determination and quantification. DIA: data-independent acquisition. (C) peptide fragmentation spectra from the spectral library. (D) Elution peak groups (left: MS1, right: MS2) from the quantification results of an example peptide AFSTNEQDKDNWNGQLK.

**Figure S2.**
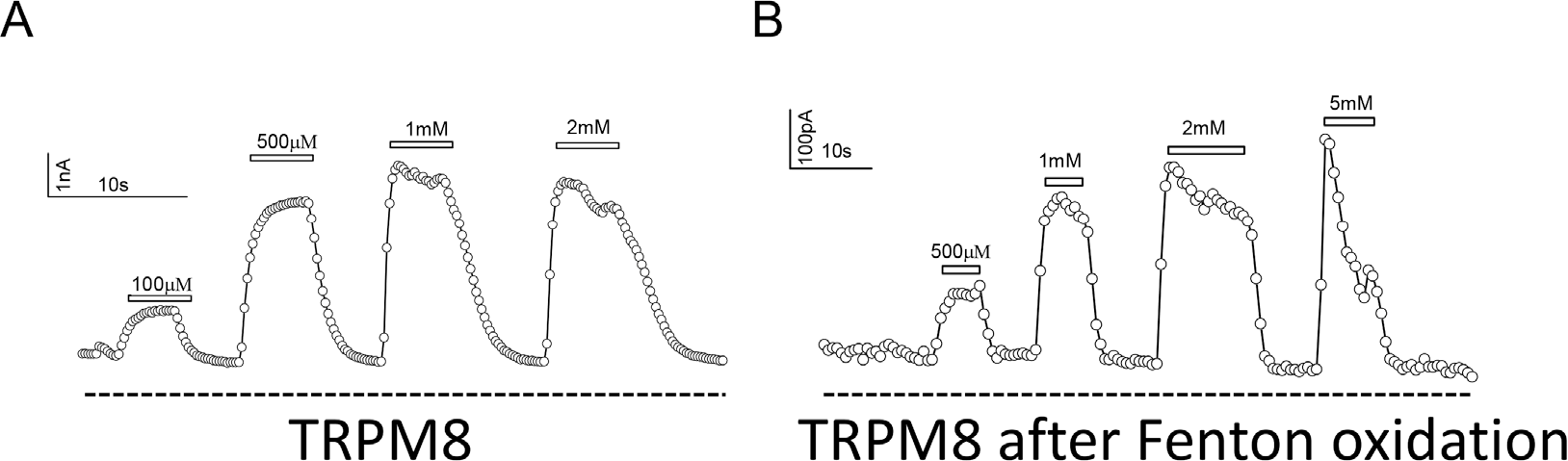
TRPM8 channel is functional after Fenton oxidation. (A and B) Representative whole-cell current recordings of TRPM8 channels activated by menthol before and after Fenton oxidation.

**Figure S3.**
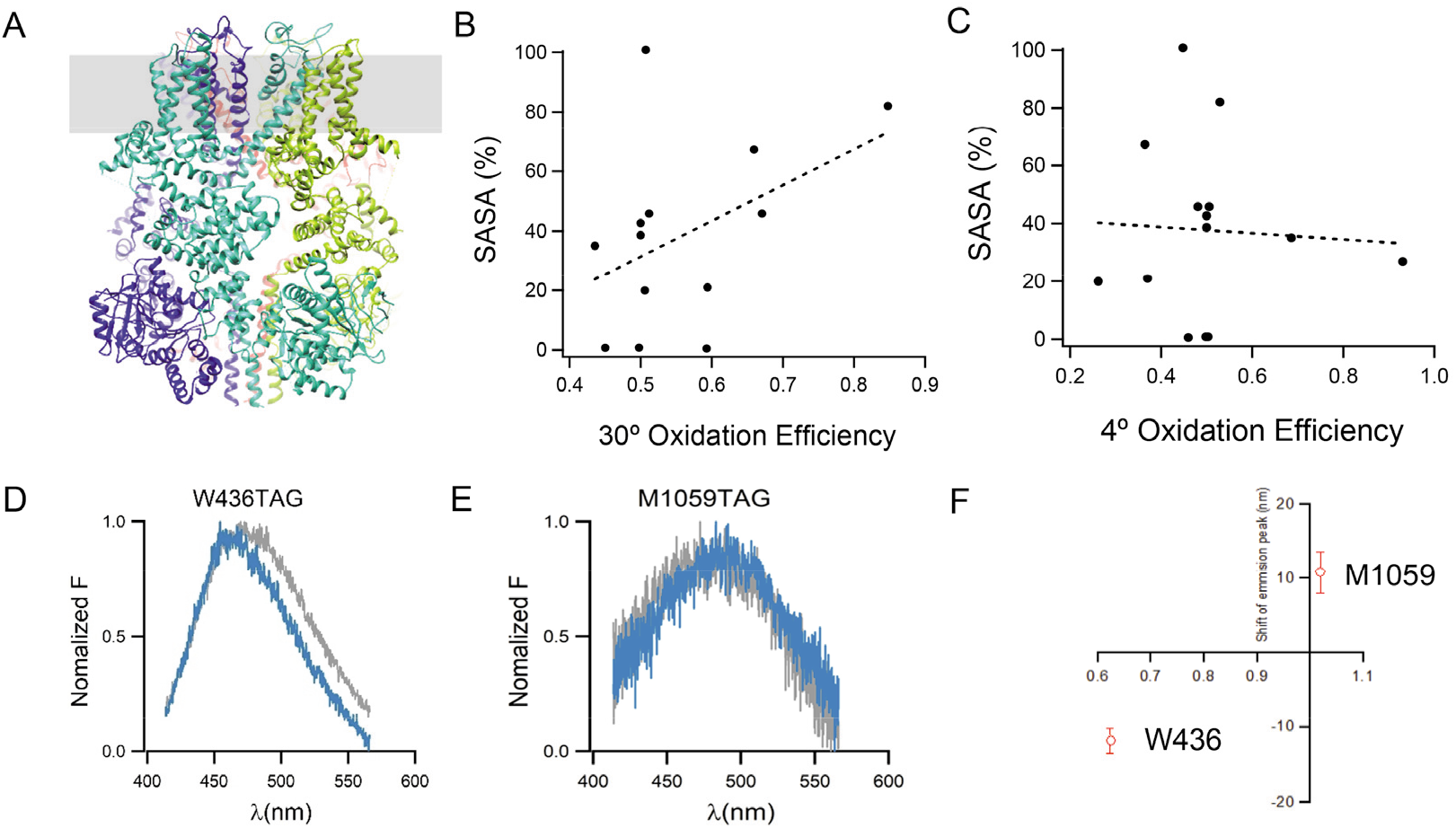
Validation of HRF-MS by cryo-EM structure determination and ANAP imaging. (A) Sideview of the cryo-EM structure of mouse TRPM8 channel in the apo and closed state. Each subunit of TRPM8 was individually colored. (B and C) SASA at the residues with buried/exposed changes found in HRF-MS was measured from the cryo-EM structure of mouse TRPM8. SASA values were plotted against the relative oxidiation efficiency determined in HRF-MS performed at either 30℃ or 4℃, respectively. (D and E) Representative emission spectra of ANAP incorporated at residue 436 and 1059, respectively. Emission spectra in grey and blue were measured at either 30℃ or 4℃, respectively. (F) Comparison of shifts in emission spectra peak of ANAP incorporated at residue W436 and M1059 with their FC values measured from HRF-MS.

**Figure S4.**
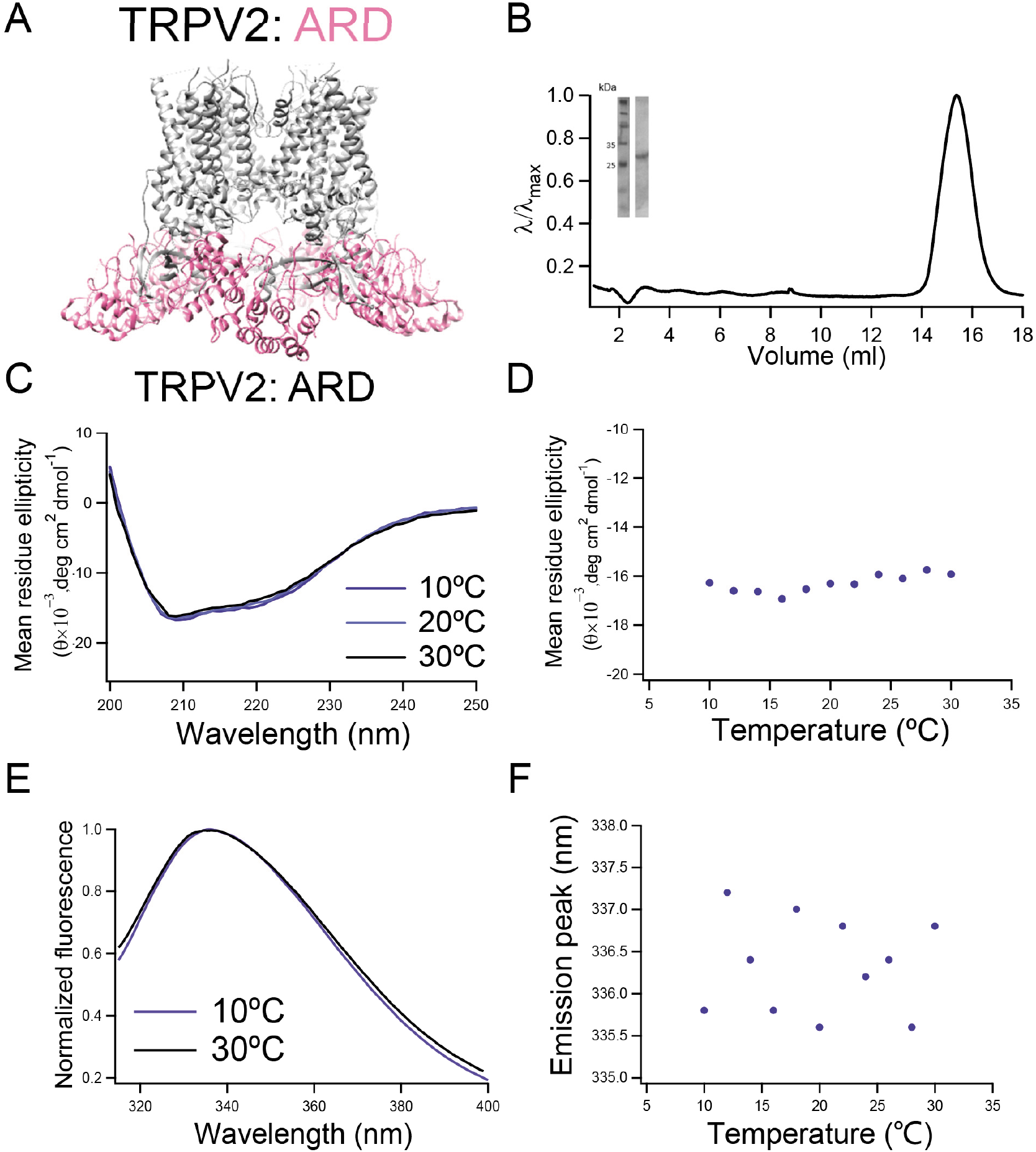
ARD in TRPV2 did not response to temperature changes from 10℃ to 30℃. (A) The location of ARD domains (colored in pink) in TRPV2. (B) Size-exclusion chromatography of the protein of ARD on Superose 6 (GE Healthcare) and SDS PAGE. (C and D) Representative CD spectra of ARD in TRPV2 and temperature dependence of CD spectra, respectively. (E and F) Representative intrinsic tryptophan emission spectra of ARD in TRPV2 and temperature dependence of intrinsic tryptophan emission peaks, respectively.

**Table S1.** Experimental summary of TRPM8 HRF-MS.

1.1 Study design for library generation.

1.2 Study design for TRPM8 oxidation experiment.

1.3 Oxidation information for TRPM8 peptides at 4°C, digested by trypsin.

1.4 Oxidation information for TRPM8 peptides at 4°C, digested by trypsin and GluC.

1.5 Oxidation information for TRPM8 peptides at 30°C, digested by trypsin.

1.6 Oxidation information for TRPM8 peptides at 30°C, digested by trypsin and GluC.

1.7 Fold changes of oxidation efficiency for TRPM8 trypsin-digested peptides at 4°C and Temperature 30°C.

1.8 Fold changes of oxidation efficiency for TRPM8 trypsin and GluC-digested peptides at 4°C and Temperature 30°C.

Relative oxidation level is calculated by Intensity(O)/(Intensity(F)+Intensity(C)). Fold changes are calculated by Relative Oxidation Level (4°C)/Relative Oxidation Level (30°C).

**Table S2.** Summary of possible oxidation sites in TRPM8 using HRF-MS.

**Table S3.** All TRPV1 mutants generated in this study and the primers for mutagenesis.

**Table S4.** All TRPM8 mutants generated in this study and the primers for mutagenesis.

### Supplementary Methods

**Commands in Rosetta to calculate SASA:**

-database /home/777/rosetta_bin_linux_2021.16.61629_bundle/main/database

-ignore_unrecognized_res

-ignore_zero_occupancy false

-load_PDB_components false

-s

/home/777/Project_TRPM8/Project_SASA/TRPV1TRPV3_PDBandSASA/7lpa.pdb

-nblist_autoupdate

-jd2:delete_old_poses

-other_pose_to_scorefile

-parser:protocol

/home/777/Project_TRPM8/Project_SASA/r_PerResidueSasaMetric.xml

-pdb_comments

-scorefile_format json

-out:output

-overwrite

### Rosetta scripts to perform SASA calculation

~~~
<ROSETTASCRIPTS>
    <SCOREFXNS>
    </SCOREFXNS>
    <RESIDUE_SELECTORS>
        <Chain name=“D” chains=“D”/>
    </RESIDUE_SELECTORS>
    <MOVE_MAP_FACTORIES>
        <MoveMapFactory name=“movemap_C” bb=“0” chi=“0”>
            <Backbone residue_selector=“D” />
            <Chi residue_selector=“D” />
        </MOVEMAPFACTORY>
    </MOVE_MAP_FACTORIES>
    <SIMPLE_METRICS>
        <SasaMetric name=“sasa” residue_selector=“D”/>
        <SasaMetric name=“sasa_polar” residue_selector=“D” sasa_metric_mode=“polar_sasa” />
        <SasaMetric name=“sasa_hydrophobic” residue_selector=”D” sasa_metric_mode=“hydrophobic_sasa” />
        <PerResidueSasaMetric name=“sasa_per” custom_type=“D” output_as_pdb_nums=“false” residue_selector=“D” mode=“all_sasa” />
    </SIMPLE_METRICS>
    <MOVERS>
        <RunSimpleMetrics name=“cache_metrics1” metrics=“sasa” prefix=“cached_total_” />
        <RunSimpleMetrics name=“cache_metrics2” metrics=“sasa_polar” prefix=“cached_polar_” />
        <RunSimpleMetrics name=“cache_metrics3” metrics=“sasa_hydrophobic” prefix=“cached_hydrophobic_” />
        <RunSimpleMetrics name=“cache_metrics4” metrics=“sasa_per” prefix=“cached_per_”/>
    </MOVERS>
    <FILTERS>

    </FILTERS>
    <PROTOCOLS>
        <Add mover=“cache_metrics1” />
        <Add mover=“cache_metrics2” />
        <Add mover=“cache_metrics3” />
        <Add mover=“cache_metrics4” />
    </PROTOCOLS>
</ROSETTASCRIPTS>
~~~

